# Complement C5 is not critical for the formation of sub-RPE deposits in *Efemp1* mutant mice

**DOI:** 10.1101/2020.12.16.423072

**Authors:** Donita L. Garland, Eric A. Pierce, Rosario Fernandez-Godino

## Abstract

The complement system plays a role in the formation of sub-retinal pigment epithelial (RPE) deposits in early stages of age-related macular degeneration (AMD). But the specific mechanisms that connect complement activation and deposit formation in AMD patients are unknown, which limits the development of efficient therapies to reduce or stop disease progression. We have previously demonstrated that C3 blockage prevents the formation of sub-RPE deposits in a mouse model of *EFEMP1*-associated macular degeneration. In this study, we have used double mutant *Efemp1*^R345W/R345W^:*C5*^-/-^ mice to investigate the role of C5 in the formation of sub-RPE deposits *in vivo* and *in vitro*. The data revealed that the genetic ablation of C5 does not eliminate the formation of sub-RPE deposits *in vivo* or *in vitro*. Contrarily, the absence of C5 in RPE cultures promotes complement dysregulation that results in increased activation of C3, which likely contributes to deposit formation even in the absence of EFEMP1-R345W mutant protein. The results also suggest that genetic ablation of C5 alters the extracellular matrix turnover through an effect on matrix metalloproteinases in RPE cell cultures. These results confirm that C3 rather than C5 could be an effective therapeutic target to treat early AMD.

## INTRODUCTION

Age-related macular degeneration (AMD) is the most common cause of visual impairment in developed countries ^1^. AMD begins as a progressive loss of fine central vision caused by degeneration of retinal pigment epithelial (RPE) cells and photoreceptors in the macular region ^1^. While some therapies can delay the progression of AMD in patients at late stages (wet AMD), there is no cure for the most common form of disease (early/dry AMD), which affects millions of people worldwide ^1^. The complex etiology of AMD and the absence of clinical biomarkers at early stages along with the diversity among patients, make it challenging to find effective therapeutic targets.

While AMD affects aged individuals, there are inherited maculopathies caused by mutations in single genes that manifest earlier in life, which makes them valuable tools to dissect the pathophysiology of AMD ^2^. For instance, *EFEMP1*-associated macular degeneration, caused by the dominant mutation p.R345W in the EGF-containing fibulin-like extracellular matrix protein 1 (*EFEMP1*) gene shares clinical features with AMD, including the early appearance of sub-RPE deposits or drusen in the macula ^3–6^. Drusen are deposits of protein and lipid that form between the basal lamina of the RPE and the connective tissue layer of the Bruch’s membrane (BrM) ^7,8^. Although the mechanisms for drusen formation and progression are not clear, it is known that the complement system, which is part of the immune system, plays an important role in drusen biogenesis ^9,10^.

The use of mouse models by our group and others has proved to be helpful for understanding the molecular mechanisms underlying the formation of sub-RPE deposits in macular degenerations, including insights about the role of the complement system in drusen formation ^4–6,11–13^. For example, we have demonstrated that mice carrying the mutation p.R345W in *Efemp1* recapitulate the formation of basal deposits underneath the RPE, the composition of which is similar to those deposits observed in patients with this mutation and in AMD patients, including increased expression of C3 in the RPE/BrM interface ^4,5^. Next, we showed that the blockage of complement activation mediated by the genetic ablation of *C3* had a protective effect, demonstrated by the absence of deposits in *Efemp1*^R345W/R345W^:*C3*^-/-^ mice ^5^. The results were recapitulated by additional *in vitro* studies using primary RPE cells from mutant mice and human ARPE19-*EFEMP1*^R345W/R345W^ cells ^5,6,14^. In addition, the mutation p.R345W in *Efemp1* resulted in decreased matrix metalloproteinase (MMP) activity, which leads to altered extracellular matrix (ECM) turnover by *Efemp1*^R345W/R345W^ RPE cells ^6,15^. Such alterations are typically observed in drusen of AMD patients as well^16^. Overall, our previous data along with observations from other labs have proven that cell-based approaches can be also used to model some aspects of AMD ^6,14,17,18^.

The complement system can be activated by three different pathways (classical, lectin, and alternative), and C3 plays a central role in all of them, so the genetic ablation of *C3* inhibits the activation of the complement system by any pathway ^19^. Activated C3b is the focus for the cleavage of C5, which initiates the formation of the terminal membrane attack complex (MAC), hence the absence of C3 prevents activation of C5 as well ^20,21^. A role for C5 in wet AMD has been demonstrated using mouse models ^13,22,23^ and some authors support the hypothesis that MAC plays a role in AMD pathogenesis, and that blocking C5 will have a protective effect for AMD patients ^24^. Yet, the blockage of C5 has not shown encouraging results in clinical trials (NTTC00935883) ^25^. Specific inhibition of C5 would prevent the generation of C5a and MAC, however, it would not prevent the complement functions associated with C3, such as immune clearance and opsonization. Indeed, the excess of C3b subunits available in a C5-depleted context would result in the formation of more C3-convertase, therefore, additional C3 activation ^26,27^. Based on our previous research, we hypothesize that downregulation of C3 rather than C5 is required for preventing the formation of basal deposits in macular degenerations ^5,6,14,17^. To address this question, in the current study, we assessed the role of C5 in the formation of sub-RPE deposits in a mouse model of the *EFEMP1*-associated macular degeneration *in vivo* and *in vitro*. Our results show that knocking out C5 is not sufficient to prevent the formation of sub-RPE deposits in *Efemp1*^R345W/R345W^ mice.

## RESULTS

### 1. *Efemp1*^R345W/R345W^: *C5*^-/-^ mice display sub-RPE deposits

Our group had previously shown a critical role for C3 in the formation of basal deposits by *Efemp1*^R345W/R345W^ mice, demonstrated by the absence of deposits in double mutant *Efemp1*^R345W/R345W^:*C3*^-/-^ mice and primary RPE cells ^5,6^. The genetic ablation of C3 also inhibits the formation of the C5-convertase and further activation of C5 and MAC, which are known to play a role in wet AMD ^22,23^. Thus, we first investigated if the inactivation of C5 was sufficient to prevent the formation of sub-RPE deposits.

Double mutant *Efemp1*^R345W/R345W^:*C5*^-/-^ mice were generated by crossing homozygous *Efemp1*^R345W/R345W^ mice and homozygous *C5*^-/-^ (B10.D2-Hc° H2^d^ H2-T18^c^/oSnJ) mice, which are serum C5 deficient ^28^. The presence of sub-RPE basal deposits was determined in 14-18 month-old mice using transmission electron microscopy (TEM). As observed in our previous study, *Efemp1*^WT/WT^:*C5*^+/+^ mice exhibit widespread basal infoldings in the RPE whereas the formation of extensive, sub-RPE basal deposits was observed in the *Efemp1*^R345W/R345W^ mice (Figure 1a, b)^4,5^. Low levels of small deposits have been observed with aging in wildtype and control mice as shown in Figure 1 (Figure 1b, c). Larger, more frequent basal deposits were present in the double mutant *Efemp1*^R345W/R345W^:*C5*^-/-^ mice (Figure 1d). The total area of basal deposits in the double mutant *Efemp1*^R345W/R345W^:*C5*^-/-^ mice trended slightly lower than in the *Efemp1*^R345W/R345W^ mice, but the differences were not statistically significant (Figure 1e, f). Thus, ablating C5 did not abolish the formation of basal deposits in *Efemp1*^R345W/R345W^:*C5*^-/-^ mice, indicating that C5 is not a good target to prevent the formation of sub-RPE deposits caused by the EFEMP1-R345W mutant protein.

**Figure 1.**
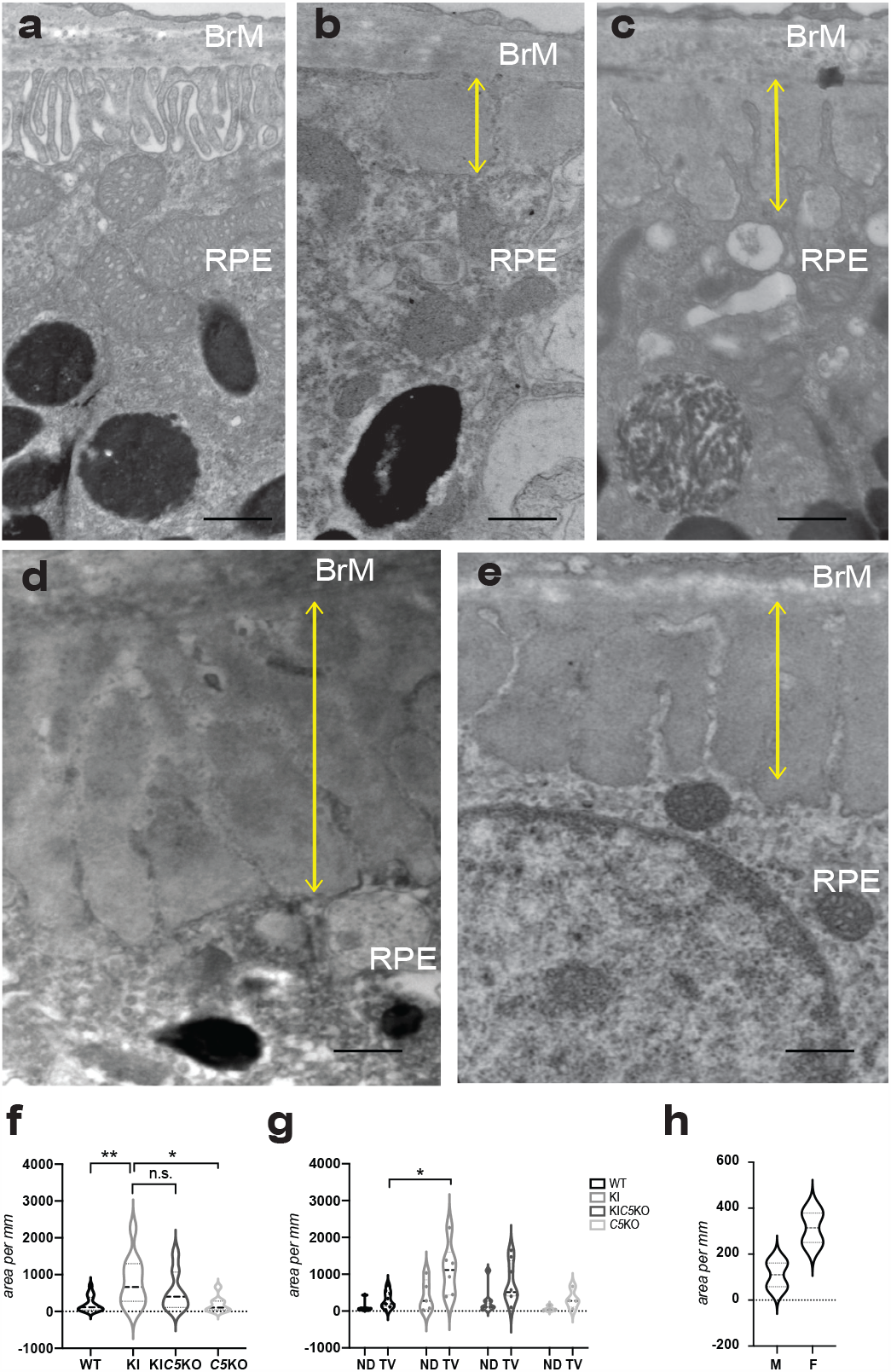
Transmission electron micrographs (TEM) of sub-RPE basal laminar deposits. **(a)** RPE with typical basal infoldings in *Efemp1*^WT/WT^:*C5*^*+/+*^ mice. **(b-e)** Basal infoldings are disrupted or missing in the regions with sub-RPE deposits in **(b)** *Efemp1*^WT/WT^:*C5*^*+/+*^ **(c)** and *Efemp1*^WT/WT^:*C5*^*-/-*^, **(d)** *Efemp1*^R345W/R345W^, and **(e)** *Efemp1*^R345W/R345W^:*C5*^-/-^ mice. Yellow lines mark the depth of the basal deposits. Scale bar = 500 nm. **(f-g)** Graphic representation of sub-RPE deposits measured as deposit area per mm length of retinal sections analyzed by TEM (average ± SD) in mice of all genotypes at 14-18 months of age. ND: nasal-dorsal, TV: temporal-ventral. **(h)** Violin plots represent the quantification of sub-RPE deposits in male vs. female *Efemp1*^R345W/R345W^ mice at 12 months as deposit area per mm of retinal section analyzed by TEM. Note that the means of cumulative areas for the deposits at 14-18 months (f-g) are substantially larger than at 12 months (g). This growth rate has been reported before. F: female, M: male. Statistical analysis by 2-way ANOVA. **p<0.01, *p<0.05, n.s. = non-significant. N=9 *Efemp1*^WT/WT^:*C5*^*+/+*^ (WT), n=6 *Efemp1*^R345W/R345W^:*C5*^+/+^ (KI), n=8 *Efemp1*^R345W/R345W^:*C5*^-/-^ (KIC5KO), n=4 *Efemp1*^WT/WT^:*C5*^-/-^ (C5KO).

Total area of basal deposits per length of the retinal slices from the temporal-ventral and nasal-dorsal quadrants at the level of the optic nerve were determined for the mutant and control mice (Figure 1g). We observed that the extent of basal deposit formation varied between retinal quadrants with deposit formation, being typically maximal in the temporal-ventral quadrant (Figure 1g). Although the morphology of basal laminar deposits was similar in both male and female mice, the amount, thickness, and continuity were greater in female mice (Figure 1h).

### 2. Primary RPE cells from *Efemp1*^R345W/R345W^: *C5*^-/-^ mice recapitulate the formation of sub-RPE deposits *in vitro*

Primary RPE cells were isolated from 2 month old mice as previously described and were cultured on transwells to maintain the RPE properties *in vitro*^6,12^. Three days post seeding, the cells were present as a confluent monolayer of hexagonal bi-nucleated cells (Figure 2a). Polarization of the RPE monolayer was demonstrated by transepithelial electrical resistance (TER) which reaches over 200 Ω·cm^2^ by 72 hours and remains stable for at least two weeks (Figure 2b).

**Figure 2.**
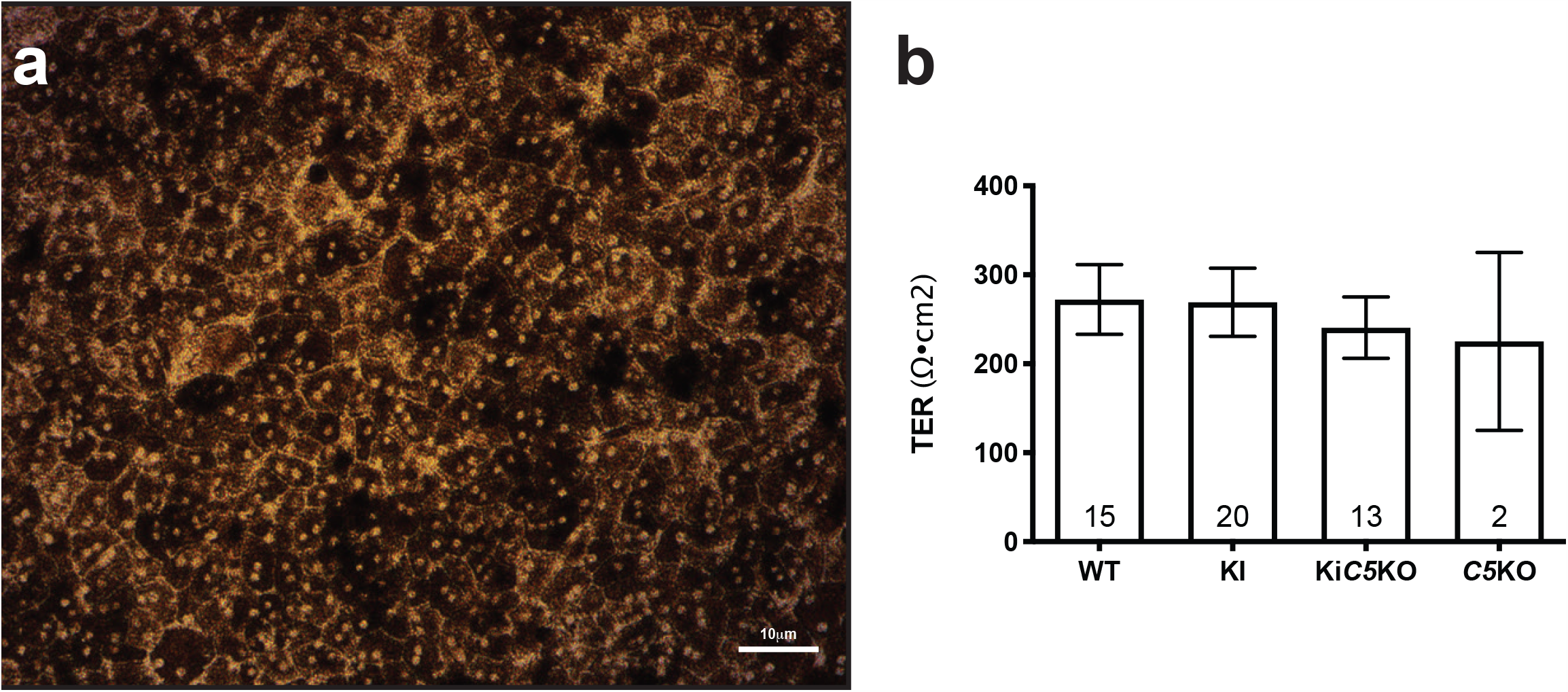
**(a)** Brightfield micrograph of mouse RPE cells *Efemp1*^R345W/R345W^:*C5*^-/-^ after two weeks on transwells confirms the formation of a healthy pigmented monolayer with honeycomb morphology. **(b)** TER measured after two weeks is similar in all cell cultures regardless the genotype.

To further investigate the role of C5 in the formation of sub-RPE deposits *in vitro*, primary RPE cells from *Efemp1*^WT/WT^:*C5*^+/+^, *Efemp1*^R345W/R345W^:*C5*^+/+^, *Efemp1*^R345W/R345W^:*C5*^-/-^, and *Efemp1*^WT/WT^:*C5*^-/-^ mice were cultured on transwells for two weeks in the absence of serum. Analysis of flat mounts of the bottom side of the insert with scanning electron microscope (SM) showed that primary cells from *Efemp1*^R345W/R345W^:*C5*^-/-^ mice made sub-RPE deposits similar to those made by the *Efemp1*^R345W/R345W^:*C5*^+/+^ cells (Figure 3), supporting that the absence of C5 is not protective against the formation of basal deposits. RPE cells from *Efemp1*^WT/WT^:*C5*^-/-^ mice were also observed to generate sub-RPE deposits, which suggests that the complement dysregulation observed *in vivo* associated with the absence of C5 is maintained by the RPE *ex vivo*, and it also results in basal deposit formation.

**Figure 3.**
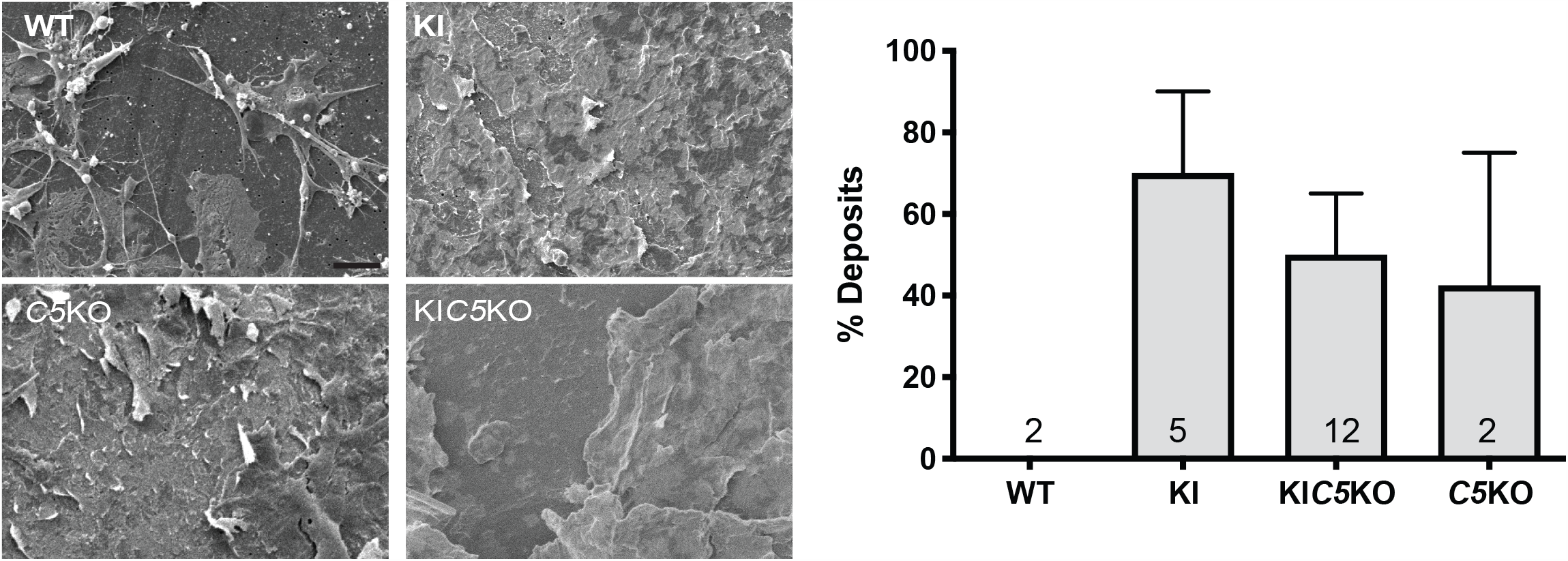
SM images of the bottom side of the transwell show filopodia that extend from the upper side through the pores of the insert in the *Efemp1*^WT/WT^:*C5*^*+/+*^ cultures (WT), vs. deposition of extracellular material in *Efemp1*^R345W/R345W^:*C5*^*+/+*^ (KI), *Efemp1*^R345W/R345W^:*C5*^*-/-*^ (KI*C5*KO) and *Efemp1*^WT/WT^:*C5*^*-/-*^ (*C5*KO) cultures. The graph represents the percentage of deposits measured in SM images using a pinpoint technique (see methods section). Data represented as average ± SEM, the numbers inside the bars indicate the amount of cultures used for quantification for each genotype. Scale bar 10 μm.

Based on our previous results using *Efemp1*^R345W/R345W^ mice ^5,6^, we characterized the RPE cell cultures and basal deposits by immunostaining of vibratome sections and flat mounts of RPE cultured on transwells. As expected, we found increased expression of EFEMP1 in *Efemp1*^R345W/R345W^:*C5*^+/+^ and *Efemp1*^R345W/R345W^:*C5*^-/-^ cell cultures compared to controls (Figure 4a)^6^. A strong immunostaining for C3 was observed in RPE cells carrying the *Efemp1*^R345W/R345W^ allele compared to WT, especially in *Efemp1*^R345W/R345W^:*C5*^-/-^ cell cultures (Figure 4a). To confirm the absence of C5 activation, we performed immunostaining with antibodies for MAC. The confocal images did not show differential activation of MAC in *Efemp1*^R345W/R345W^:*C5*^+/+^ mutant cells compared to *Efemp1*^WT/WT^:*C5*^+/+^ (Figure 4a). As expected, we did not detect MAC in *C5*^-/-^ cells (Figure 4a).

**Figure 4.**
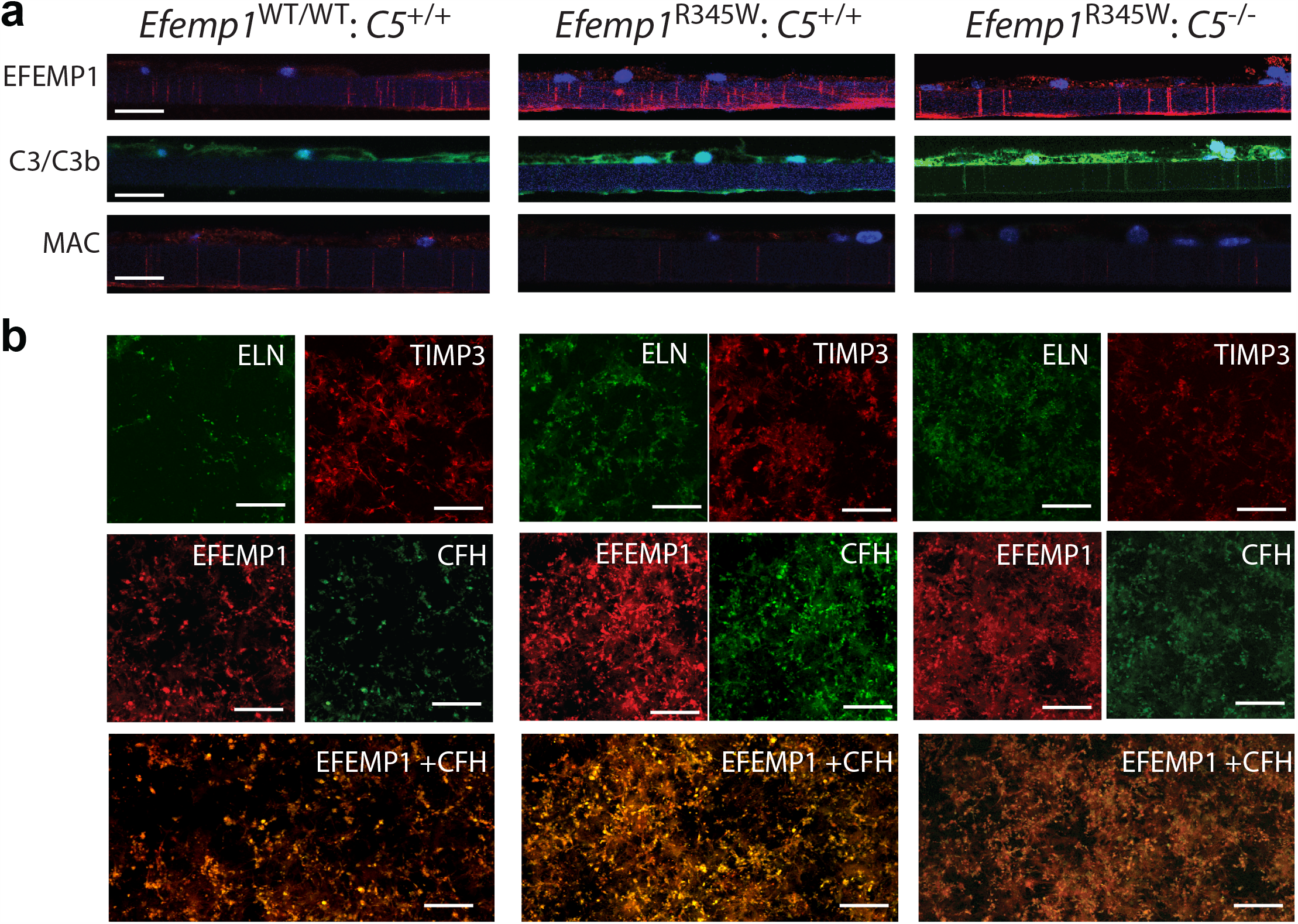
**(a)** Vibratome sections of primary mouse RPE cells *Efemp1*^WT/WT^:*C5*^*+/+*^, *Efemp1*^R345W/R345W^:*C5*^*+/+*^, and *Efemp1*^R345W/R345W^:*C5*^*-/-*^ cultured on transwells for two weeks and immunostained with antibodies for EFEMP1, C3/C3b, MAC. Note positive C3 staining at the bottom of the insert in *Efemp1*^R345W/R345W^:*C5*^*+/+*^ cultures and throughout the insert pores in *Efemp1*^R345W/R345W^:*C5*^*-/-*^ cultures. This is due to the orientation of the sections. **(b)** Flat mounts of the bottom side of the inserts were immunostained with antibodies for ELN, TIMP-3, EFEMP1, CFH. EFEMP1 and CFH colocalize in the deposits. Scale bars a-i: 25 μm, j-aa: 100 μm.

To characterize the composition of the sub-RPE deposits made by primary cells *in vitro*, flat mounts of the bottom side of the inserts containing RPE cells were immunostained with antibodies for the ECM proteins that typically accumulate in basal deposits of AMD patients ^29^. As previously reported for *Efemp1*^R345W/R345W^ mutant mice and RPE cultures, we detected accumulation of elastin, TIMP3, and EFEMP1 secreted by the RPE to the bottom side of the insert in *Efemp1*^R345W/R345W^:*C5*^-/-^ cultures (Figure 4b) ^29^. In this study, EFEMP1 was homogeneously deposited along the insert, and accumulated in *Efemp1*^R345W/R345W^:*C5*^+/+^ and *Efemp1*^R345W/R345W^:*C5*^-/-^ cultures compared to wildtype. Excessive deposition of CFH associated with the presence of EFEMP1-R345W mutant protein was observed in *Efemp1*^R345W/R345W^:*C5*^+/+^ and *Efemp1*^R345W/R345W^:*C5*^-/-^ cultures (Figure 4b). Of note, EFEMP1 and CFH co-localized on the deposits (Figure 4b), which confirms the interaction of these two proteins in the ECM ^30^.

### 3. RPE cells from *Efemp1*^R345W/R345W^: *C5*^-/-^ mice secrete increased amounts of C3

We have previously demonstrated that C3 is necessary for the formation of basal deposits in mice and RPE cells with the p.R345W mutation in *Efemp1* ^5,6^. Our previous experiments using primary mouse RPE cells showed that *Efemp1*^R345W/R345W^ cells secreted increased amounts of C3 ^6^. In the current studies, we measured the amount of C3 secreted to the conditioned media by wildtype and mutant cells via ELISA. In concordance with the strong C3 immunostaining observed in *Efemp1*^R345W/R345W^: *C5*^-/-^ cells, elevated levels of C3 were detected in conditioned media from *Efemp1*^R345W/R345W^: *C5*^+/+^ cells (2-way ANOVA, p=0.0195) compared to wildtype (Figure 5a). In the absence of C5, the levels of C3 detected in conditioned media from *Efemp1*^R345W/R345W^: *C5*^-/-^ cells increased an additional 50% (2-way ANOVA, p=0.0035). Despite a large variability among samples, transcription levels of *C3* were also higher in the context of the *Efemp1*^R345W^ allele, although not significantly increased in *C5*^-/-^ cells (Figure 5b). Based on these results, amplified complement activity in this system arises from increased posttranslational activation of C3.

**Figure 5.**
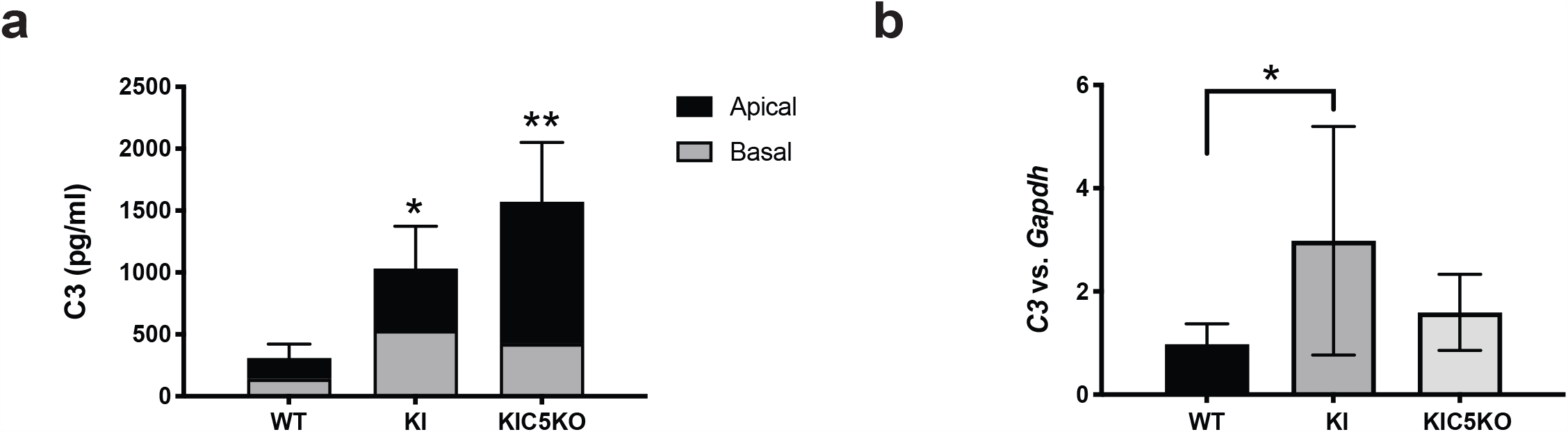
**(a)** Levels of C3 measured by ELISA in the apical and basal conditioned media of RPE cultures showed increased levels of C3 in *Efemp1*^R345W/R345W^:*C5*^*+/+*^ and *Efemp1*^R345W/R345W^:*C5*^*-/-*^ cultures compared to controls. Data represented as average ± SD (2-way ANOVA, *p<0.05, **p<0.01). **(b)** mRNA levels of *C3* measured in RPE cells cultured for two weeks on transwells in the absence of serum, and normalized to *GAPDH* showed significant increase of C3 expression. Data represented as average ± SD (unpaired t-test, **p<0.01).

### 4. Depletion of C5 restores MMP-2 activity in mutant *Efemp1*^R345W/R345W^ RPE cells

We have previously reported that MMP-2 activity diminishes in conditioned media of mouse *Efemp1*^R345W/R345W^ RPE cells, and it does not change in the presence or absence of C3^6^. To assess whether the absence of C5 had an impact on the ECM turnover, we used zymography analyses, which allows measurement of the collagenase activity of MMP-2, which is secreted by the RPE and mediate the ECM turnover of the BrM^31^. The collagenase activity detected in conditioned media of *Efemp1*^R345W/R345W^: *C5*^-/-^ cells was similar to the activity found in wildtype controls, which differs from the MMP-2 activity found in *Efemp1*^R345W/R345W^: *C5*^+/+^ and *Efemp1*^R345W/R345W^: *C3*^-/-^ cells (Figure 6). Hence, the absence of C5 restores the alterations in ECM turnover associated with the EFEMP1 mutant protein.

**Figure 6.**
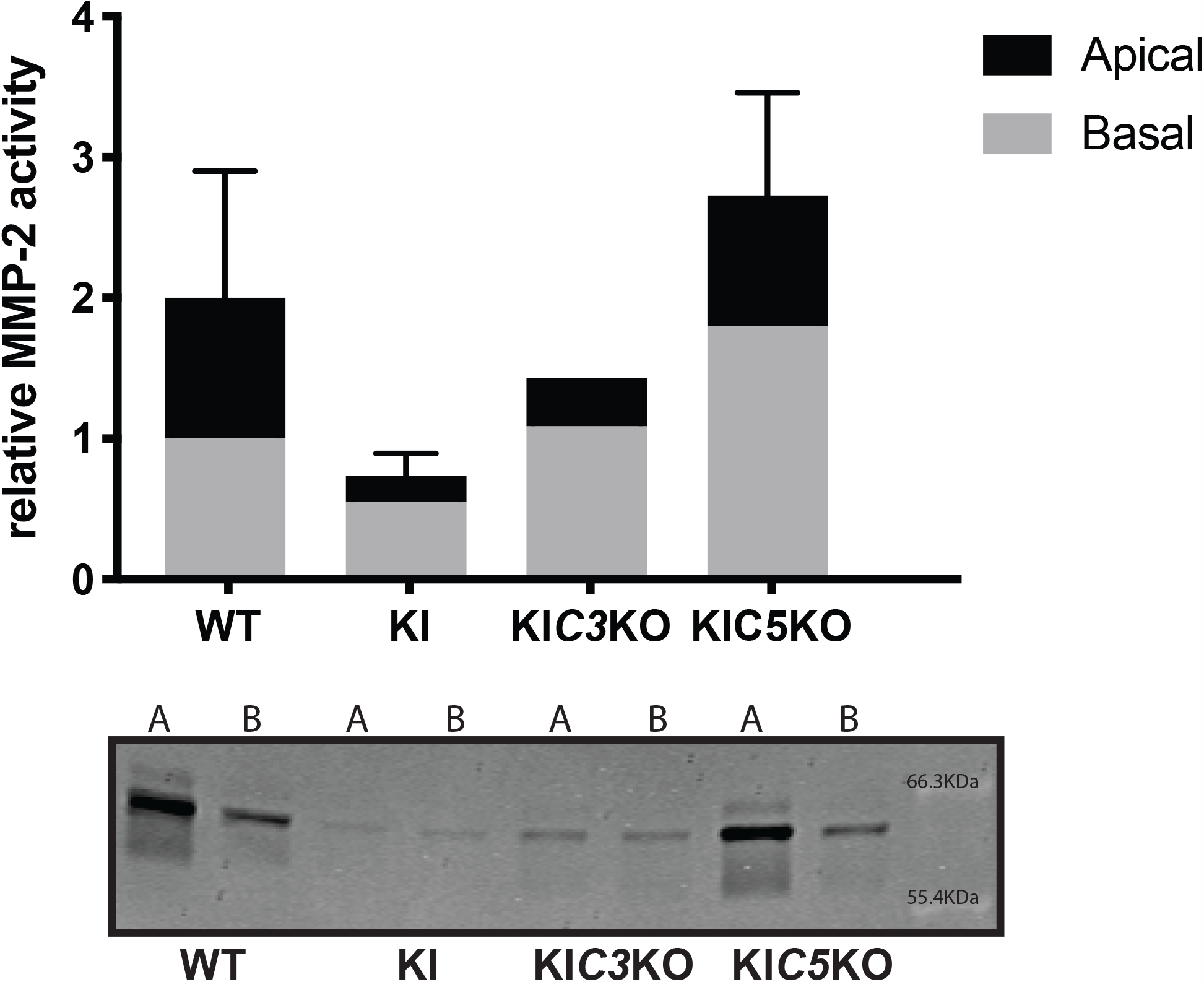
MMP-2 activity measured by zymography in apical and basal conditioned media of *Efemp1*^WT/WT^:*C5*^*+/+*^, *Efemp1*^R345W/R345W^:*C5*^*+/+*^, *Efemp1*^R345W/R345W^:*C3*^*-/-*^, and *Efemp1*^R345W/R345W^:*C5*^*-/-*^ cultures demonstrated that the ECM turnover is altered only in the absence of C3, but it remains normal in the absence of C5.

## DISCUSSION

Genetic and clinical studies have demonstrated a role for the complement system in the formation of drusen in early stages of AMD, but defining the extent to which each complement component contributes to this disease is critical for the identification of efficient therapeutic targets ^29,32–38^. To address this need, in this study we have used mouse and cell-based models of macular degeneration to investigate the role of C5 in the formation of sub-RPE deposits ^4–6,17^. The *in vivo* and *in vitro* studies showed that the genetic ablation of C5 does not protect against the formation of basal deposits underneath the RPE in a mouse model of the inherited *EFEMP1*-associated macular degeneration. Indeed, the ablation of C5 results in accumulation of basal deposits underneath the RPE *in vivo* and *in vitro* even in the absence of the EFEMP1-R345W mutant protein. In addition, blockage of C5 rescues the MMP activity disrupted by the mutation p.R345W in *Efemp1*, however, it enhances the activation of the complement system mediated by increased amounts of C3 secreted by the RPE in culture. Consistent with previous studies in our lab, the increased levels of C3 observed in *Efemp1*^R345W/R345W^: *C5*^-/-^ cell cultures point at C3 as a major contributor for basal deposit formation ^5,6,14,17^. These results could explain the failure of C5-targeted treatments in clinical trials and supports the potential of C3 as a therapeutic target to reduce drusen progression.

For the first time, we have characterized the regional localization of sub-RPE deposits in *Efemp1*^R345W/R345W^ mice, which are more abundant in the temporal-ventral quadrant. This finding could be explained by the higher RPE cell density, including more bi-nucleated cells, present in the ventral-central region of mice retinas^39^. Likewise, in humans, RPE cell density is highest in the central temporal retina^40^, where the macula is located, and where the rate of apoptosis increases with age^41^. Higher levels of deposit formation were observed in female mice, which could be explained by a deficiency of estrogen described in older female rodents^42^. In humans, the beneficial effect of estrogens for the retina has been demonstrated by a higher risk of AMD in women who enter menopause at young age^43^. In mice, age-related estrogen deficiency increases susceptibility to sub-RPE deposition caused by dysregulating turnover of BrM, which contributes to thickening of collagenous layers ^44^. Other studies in our lab have demonstrated a key role for the BrM turnover in the formation of sub-RPE deposits ^6,14,15^. Also, estrogen can regulate signaling pathways involved in AMD pathogenesis through its anti-oxidative and anti-inflammatory effect ^45^. Indeed, gender specific effects on autoimmune disorders and on immune cell populations are beginning to be appreciated ^46^. Further studies should be performed to determine the protective role of estrogens in the retina.

In the absence of complement C5, the number and size of basal laminar deposits in *Efemp1*^R345W/R345W^ mice were diminished but not eliminated. This is in contrast to previously reported results that demonstrated that genetic ablation of C3 prevented the formation of basal laminar deposits due to the EFEMP1-R345W mutant protein^5,6^. Therefore, inhibition of C5 is not protective against deposit formation. Indeed, the absence of C5 is sufficient to propel accumulation of material underneath the RPE *in vivo* and *in vitro*, likely associated with the positive feedback loop that increases C3^27^. This dysregulation of the complement system caused by a long-term C5 inhibition is typically observed in patients with glomerular disease, which develop C3-associated deposits in the kidney comparable to those developed by patients with deficiencies in CFH and CFI ^27,47^. Our *in vitro* model offers a pure environment without external complement regulators normally found in serum and tissues. In such system, the high density of C3b will lead to the formation of more convertase, which has great affinity for C3 and may cleave it in the absence of a better substrate like C5, generating the positive feedback loop measured as increased levels of C3 in RPE cells and conditioned media of *Efemp1*^R345W/R345W^: *C5*^-/-^ cultures ^21,26^.

As we reported previously, the sub-RPE deposits found in primary cultures of both *Efemp1*^R345W/R345W^: *C5*^+/+^ and *Efemp1*^R345W/R345W^: *C5*^-/-^ mice are comprised of excessive depositions of normal ECM proteins^6,14^. Positive immunostaining with antibodies for EFEMP1, TIMP-3 and elastin, confirmed that the deposits made by RPE cells *Efemp1*^R345W/R345W^: *C5*^+/+^ and *Efemp1*^R345W/R345W^: *C5*^-/-^ are similar in composition to basal laminar deposits and drusen found in AMD patients^29^. The immunostaining also revealed an intensified deposition of CFH underneath the RPE in the presence of EFEMP1-R345W mutant protein. This phenomenon can be explained by the fact that EFEMP1 binds CFH^30^. Wyatt and collaborators demonstrated a higher affinity of the AMD-risk associated variant CFH-402H for EFEMP1 compared to the low-risk allele CFH-402Y^30^. Given that the mutation p.R345W is located in the protein domain where EFEMP1 binds CFH, it is possible that mutant EFEMP1-R345W has higher affinity for CFH than EFEMP1-WT, which would favor the inclusion of CFH in the protein aggregates typically originated by EFEMP1-R345W^6^. CFH sequestered into protein aggregates will likely lose its complement regulatory function, which contributes to the increased levels of C3 found in these cultures. This process shares features with the impaired complement regulation in the BrM with age, associated with the loss of heparan sulfate proteoglycan, which is responsible for anchoring CFH to the BrM, and that is exacerbated in the context of the AMD risk allele *CFH*-Y402H^48^.

Based on our previous studies, we expected the MMP-2 activity to be diminished in the conditioned media of *Efemp1*^R345W/R345W^ RPE cells ^6^, however, in the absence of C5, the MMP-2 activity was restored to normal levels, which would reinstate the normal ECM turnover. We and others have demonstrated that C3a can modulate the activity of MMP-2^6,17,49^. Thereby, the increased MMP-2 activity observed in the *Efemp1*^R345W/R345W^:C5^-/-^ cultures could be a secondary effect driven by overactivation of C3.

Based on this and other studies in our lab, we believe that the increased activation of C3 in *Efemp1*^R345W/R345W^ cells occurs via tick-over, through deposition of C3b on the EFEMP1-derived abnormal ECM, which prevents its futile depletion and enhances the stabilization of the tick-over convertase on the substrate ^6,14,50^. This hypothesis is sustained by the positive immunostaining for C3b in the abnormal ECM secreted by mutant *Efemp1*^R345W/R345W^ RPE cells. Of note, C3b deposited on the ECM that is not inactivated by CFH will result in chronic activation of C3 by the mediated by the canonical C3-convertase of the alternative pathway, which is also dysregulated in the absence of C5^21,50,51^. In any case, complement activity would be mediated by C3, but does not require the activation of C5 or MAC. Further analyses would be required to define whether the activation of C3 occurs via tick-over or the canonical pathway.

In conclusion, blockage of C5 is not sufficient to protect against the formation of sub-RPE deposits, which may explain the inefficiency of anti-C5 therapies in the clinic^25,52^. These findings suggest that therapeutic strategies to prevent C3 rather than C5 activation will prove more effective to treat patients with early-intermediate stage AMD.

## METHODS

### Mice

The guidelines of the ARVO Statement for the Use of Animals in Ophthalmic and Vision Research were followed. *Efemp1*^R345W/R345W^ mice, made and characterized previously, were generated from het x het crossings^4^. Wildtype littermates generated during het x het crosses of *Efemp1*^WT/R345W^ mice were used as controls. *Efemp1*^R345W/R345W^ mice were crossed with homozygous C5^-/-^ mice (B10.D2-Hc^0^ H2^d^ H2-T18^c^/o2SnJ) purchased from Jackson Laboratory (JAX stock #000461), which have a 2 base deletion that creates a STOP codon and results in a non-functional protein. None of these mice contained the *rd8* mutation^53^. Mice were genotyped for the *Efemp1* mutation as previously described^5^. For the C5 genotype, DNA was amplified by PCR using the following primers: forward 5’ TTGCTTCCACAGGTATGGTG 3’ and reverse 5’ CCCCACCCTCTTCTGGTACT 3’ and sequenced by Sanger.

### Assessment of basal deposit formation by TEM

At 14-18 months mice were euthanized and immediately ‘perfused’ with 4% paraformaldehyde. Before removing an eye, its orientation was marked using a fine cautery tip. The eyes were promptly placed in half strength Karnovsky’s fixative (2% formaldehyde + 2.5% glutaraldehyde, in 0.1 M sodium cacodylate buffer, pH 7.4). After 4 hours, incisions were made at the level of the *ora serrata* and the cornea and lens were removed. The eye cups were kept in the fixative overnight at 4° C, rinsed three times in 0.1M sodium cacodylate buffer and stored at 4° C until they were embedded. The post-fixation, *en bloc* staining, embedding, sectioning of the blocks and assistance with the imaging were all performed in the Schepens/MEE Morphology Core. Samples were post-fixed with 2% osmium tetroxide in 0.1M sodium cacodylate buffer for 1.5 hours, *en bloc* stained with 2% aqueous uranyl acetate for 30 minutes, and then dehydrated with graded ethyl alcohol solutions. The samples were resin infiltrated in ethyl alcohol and Spurr’s resin epoxy mixtures using an automated EMS Lynx 1 EM tissue processor (Electron Microscopy Sciences). Processed samples were infiltrated with two changes of fresh Spurr’s resin for 24 hours and the samples were then oriented in the resin. The resin was polymerized within silicone molds using an oven at 60°C for a minimum of 24 hours. The samples were en bloc stained with 2% uranyl acetate for 30 minutes. Semi-thin sections for light microscopy were cut in cross-section at 1-micron and stained with 1% toluidine blue in 1% sodium tetraborate aqueous solution for assessment and screening regions of the processed samples for thin sectioning. Ultrathin sections (70 nm) were cut using a diamond knife and were collected onto single slot formvar-carbon coated grids. The retinal sections were cut at the level of the optic nerve and extended from the optic nerve outwards to the ora serrate in the temporal/ventral quadrant and in the dorsal/nasal quadrant. The sections were not post stained. The grids were imaged at a direct magnification of 18,500 using a FEI Tecnai G2 Spirit transmission electron microscope at 80 kV. The microscope was interfaced with an AMT XR41 digital CCD camera for digital TIFF file image acquisition.

To systematically assess basal laminar deposit formation, images were acquired of all basal deposits along the entire length of each thin section at a direct magnification of 18,500. Both the temporal/ventral and dorsal/nasal quadrants at the level of the optic nerve were analyzed since deposit formation was not equivalent within the 4 retinal quadrants. The areas of the deposits were determined by manually outlining the deposits in each image and determining the areas using ImageJ. If the images were not acquired at a direct magnification of 18,500 the results were normalized to that magnification. The areas of all deposits were summed for the sections from the two quadrants per mouse and reported as area (µ^2^/mm). Deposits were present in the peripapillary region of the sections of most eyes. The areas of these deposits were excluded from the analyses because they were associated with incompletely formed or degrading RPE cells. For all samples in which the deposits in these areas were measured, the areas were similar in each eye. A minimum of 65 micrographs were taken per mouse.

### Mouse RPE cell isolation

RPE cells were harvested from 10-week-old mice following CO2-induced euthanasia. Eyes were dissected and RPE cells were collected by enzymatic digestion as previously described^6,12^. Briefly, eyecups were dissected and the neural retina was separated from the RPE with 1 mg/ml of hyaluronidase. The RPE cells were detached from the BrM with trypsin and collected in RPE media with 20% FBS. Isolated cells were resuspended in RPE media containing 5% of FBS and seeded on 6.5mm transwells coated with laminin following published methods^6,12^.

### RPE cell cultures

Cells were cultured at 37°C in 5% CO2 under a humidified atmosphere changing media twice a week. Serum was removed at least 72 hours before performing any experiment. Transepithelial electrical resistance (TER) of the RPE cultures were measured using an epithelial voltohmmeter (EVOM)^54^.

### RNA and qRT-PCR

After 2 weeks in culture, RPE cells were lysed with RLT buffer containing b-mercaptoethanol, and RNA was extracted with DNA/RNA/Protein Mini Kit (Qiagen). The quality and quantity of RNA was assessed using the Agilent RNA 6000 Nano Kit Bioanalyzer (Agilent Technologies). All samples had RIN values between 9 and 10. mRNA expression levels were analyzed by qRT-PCR and normalized to *Gapdh* as an endogenous reference. cDNA was synthesized using Affinity Script cDNA Synthesis kit (Agilent Technologies). 5 ng of cDNA, 200 nM of each primer and 10 µl of Fast SYBR Green were combined. Amplification was done in the Stratagene Mx3000P® QPCR system using the following program: 95°C for 20 s, 40 cycles of 95°C for 3 s, 60°C for 30 s followed by melting curve. Each sample was assayed in triplicate. For each experiment the expression level of the sample from the wildtype mouse was set to a value of 1. Primers used for amplification are the following: *C3* (forward *5’-*TCCTGAACTGGTCAACATGG-3’ and reverse 5’-AAACTGGGCAGCACGTATTC-3’).

### ELISA

Apical and basal conditioned media from RPE cell cultures were collected after 2 weeks in culture and concentrated to equal volumes through 10 kDa Amicon filters (Millipore, Billerica, MA). The fraction over 10 kDa was used to quantify C3 using the ALPCO ELISA kit.

### Characterization of Deposits in vitro

Deposits were characterized by SM and immunofluorescence as previously described^6^.

### Scanning electron microscopy

RPE cells grown on inserts were fixed in cold 4% PFA in PBS followed by fixation in 1% glutaraldehyde, washed in PBS and then in dH2O and dehydrated by serial ethanols, 35%, 50%, 70%, 95%, 95% and 100% followed by critical dehydration using the SAMDRI-795 system. After dehydration, inserts were split into 2 pieces and the top or bottom side was coated with chromium using a Gatan Ion Beam Coater for 10 min. Coated inserts were imaged by Field Emission Scanning Electron Microscope (JEOL 7401F).

The percentage of deposit formation was defined as the percentage of SM images positive for deposits divided by the total number of images taken of the whole insert (# images per equal parts insert). A minimum of 10 images was taken per sample. The experimenter was masked to the sample identity.

### Vibratome sections

RPE cells on transwells were fixed for 10 min in cold 4% paraformaldehyde (PFA) in PBS followed by fixation in 1% glutaraldehyde for 30 min at RT. The insert was then removed and split with a razor blade into small fragments of 2.5 mm x 5 mm, which were embedded into 10% Agarose XI as previously published^12^. Sections of 100µm were performed at medium speed (around 5) and amplitude (approx. 6) using a vibratome Leica VT1000S (Leica). Sections were placed on a glass slide for immunostaining.

### Immunostaining

Cell inserts were rinsed in PBS, fixed for 10 min in cold 4% PFA in PBS followed by fixation in 1% glutaraldehyde for 30 min at RT. The inserts were cut off with a razor blade and stored in PBS at 4°C pending sectioning and immunohistochemical analyses. To analyze the bottom side of the inserts, they were placed face down on slides as flat mounts. For staining, slides containing flat mounts or vibratome sections were blocked with 1% BSA and incubated with primary antibodies anti C3/C3b (AB11862, Abcam, Cambridge, MA), EFEMP1 (SDIX), MAC (C5b-9 AB55811, Abcam, Cambridge, MA), TIMP-3 (AB39184 Abcam, Cambridge, MA), ELN (sc17581, Santa Cruz Biotechnology, Santa Cruz, CA), and CFH (AB53800, Abcam, Cambridge, MA) overnight at 4°C. Secondary antibodies labeled with Alexa-488 or Alexa-555 were incubated for 1 h at RT. Slides were mounted with Fluoromount G and visualized by TCS SP5 II confocal laser scanning microscope (Leica). Samples incubated with 1% BSA instead of primary antibody were used as negative controls.

### MMP-2 activity

was measured in conditioned media from RPE cultures by zymography as previously described^6^. Briefly, 5 µl of equal volume supernatants were loaded onto Novex 10% gelatin gels (Life Technologies, Grand Island, NY). Zymography assays were then performed per manufacturer’s instructions. Gels were scanned using the Odyssey system (Li-Cor, Lincoln, NE). MMP-2 was identified by molecular weight. Gelatinase activity was quantified using densitometry and the software ImageStudioLite (Li-Cor, Lincoln, NE).

### Statistical analyses

Results are expressed as mean ±SD or SEM, with p<0.05 considered statistically significant. Differences between groups were compared using the Student t-test or ANOVA as appropriate using GraphPad Prism software.

## ACKNOWLEDGEMENTS

This work was supported by the Ocular Genomics Institute. The TEM work was supported in part by a National Eye Institute Core grant (P30EY003790).

The authors would like to thank Schepens Eye Research Institute morphology core facility, especially Philip Seifert for performing the TEM, and Ann Tisdale for helpful technical assistance with SM.

## AUTHOR CONTRIBUTIONS

D.L.G. performed the *in vivo* experiments and helped writing the manuscript. E.A.P. supervised the work and edited the manuscript. R.F.G. bred and genotyped the mice, performed all the *in vitro* experiments and wrote the manuscript.

## CONFLICT OF INTEREST STATEMENT

The authors have declared that they have no conflicts of interest.

## Notes

### Competing Interest Statement

The authors have declared no competing interest.

